# Initial efficacy determination and resistance profile of anti-*Acinetobacter* antibiotics, turnercyclamycins

**DOI:** 10.1101/2021.11.01.466777

**Authors:** Bailey W. Miller, Albebson L. Lim, Margo G. Haygood, Eric W. Schmidt

## Abstract

Drug-resistant *Acinetobacter* is a challenging, deadly pathogen of increasing prevalence in the US healthcare system. Recently, we described a series of lipopeptides, the turnercyclamycins, which retain potency against *Acinetobacter* strains that are resistant to the last-line antibiotic, colistin. To further evaluate the potential of turnercyclamycins, we completed mouse efficacy, pharmacokinetics, and toxicity studies. These demonstrate that turnercyclamycin A has a pharmacological profile with similarity to other lipopeptides that are in clinical use. Turnercyclamycin A was well tolerated in mice up to 25 mg/kg, and exhibited >99% and >98% reduction in bacterial load compared to vehicle control in a thigh infection model at 25 and 12.5 mg/kg, respectively. This result closely reflected the anticipated effectiveness based upon *in vitro* activity and was similar to the colistin control. *Acinetobacter* strains resistant to colistin often harbor the *mcr-1* resistance gene. Here, we show that the effectiveness of turnercyclamycins against *Escherichia coli* is not greatly altered by *mcr-1* (0- to 2-fold) whereas there is a 16-fold increase in the colistin minimal inhibitory concentration when *mcr-1* is present. These data suggest that turnercyclamycins are suitable for further investigation and optimization as anti-*Acinetobacter* lead compounds.

## Introduction

Carbapenem-resistant *Acinetobacter baumannii* is characterized as an urgent threat by the CDC. The 2019 Antibiotic Resistance Threats report estimated 8,500 infections occur in hospitalized patients annually in the US, resulting in an estimated 700 deaths and over 280 million dollars in healthcare costs^1^. *Acinetobacter* is particularly adept at acquiring resistance mechanisms, and there has been a rapid rise in clinical isolates with resistance to almost every class of clinically used antibiotics.^2^ The last line antibiotic that remains for the treatment of these multidrug-resistant infections is colistin,^3,4^ however strains with colistin resistance are also increasingly on the rise^5^. One contributing factor to this rise is attributed to the recently discovered mobile colistin resistance gene, *mcr-1*, which is a plasmid-born resistance marker that is capable of horizontal transfer. The growing threat of *Acinetobacter* infections and rapidly developing resistance profiles underline the urgent need for new therapeutics that circumvent *Acinetobacter* drug resistance and can act as new therapeutics for Gram-negative pathogens.

The turnercyclamycins were isolated from *Teredinibacter turnerae*, an intracellular symbiont of shipworms, which are wood-consuming marine bivalve mollusks. These cyclic lipodepsipeptide antibiotics are the products of an NRPS gene cluster and were targeted for isolation based on genomic and metagenomic analysis of isolates and shipworm tissues. They are reasonably potent agents against *Acinetobacter baumannii* and *E. coli*, with minimum inhibitor concentrations of 8 and 1 mcg/mL, respectively. Turnercyclamycins retain their activity against clinical *Acinetobacter* complex strains that are resistant to colistin, the last-line clinical agent for *A. baumannii* infections. While it was shown that these compounds similarly disrupt the outer membrane of Gram-negative pathogens, the exact molecular mechanisms appear to be distinct in the two molecular families.^6^

In order to assess the pharmacological potential of these molecules, a series of *in vitro* resistance and *in vivo* pharmacokinetic (PK), toxicology, and efficacy studies were completed. First, the mcr-1 resistance cassette, which is a growing threat to antibiotic susceptibility globally, was transformed into *E. coli*. Encouragingly, turnercyclamycin susceptibility was not affected by mcr-1, while the MIC for colistin was increased 16-fold. In murine models, turnercyclamycins demonstrate a similar PK profile to known lipopeptide antibiotics that are currently in use in the clinic. They are also well tolerated in mice at doses that show statistically significant efficacy in murine thigh infection models with *Acinetobacter baumannii*. Taken together, these data reinforce the potential utility of turnercyclamycins as anti-*Acinetobacter* agents in the face of rising global antibiotic resistance.

## Results

### *in vivo* PK, toxicity, and efficacy studies

To investigate the pharmacokinetics, toxicity, and efficacy of turnercyclamycins, we selected turnercyclamycin A (**1**), which is isolated in higher yield and is more easily purified than other naturally occurring analogs. The producing strain *T. turnerae* T7901 was fermented, and turnercyclamycin A was rigorously purified to >99% purity using diode array HPLC and quantified using a precision analytical balance. The purified compound was shipped to Eurofins (Taiwan), which performed the studies. All studies were performed using IV injection with **1** formulated in 3% ethanol/ phosphate buffered saline (PBS).

The experimental design is summarized in **Table S1**. Briefly, twenty-four immune competent, uninfected female BALB/c mice were used to assess the pharmacokinetic characteristics of **1. 1** was IV administered by slow bolus (BID) at 5 mg/kg, and plasma samples were collected at 0.05, 0.165, 0.5, 1, 2, 4, 8, and 24 hour(s) post-infusion. Plasma concentrations of **1** were determined at each time point by LC-MS/MS (**Figure S1**), and fundamental PK parameters were obtained through non-compartmental analysis (NCA) using WinNonlin (**Table 1**). A clearance of 17.4 mL/hr/kg and Vss of 0.1 L/kg led to a half-life of 4.35 hours. These values are comparable to other lipopeptides. For example, daptomycin (a lipopeptide that is Gram-positive selective) exhibited a similar low volume of distribution (0.1 L/kg) a half-life of ∼8 hours in human volunteers. This half-life and clearance rate indicates that twice daily (bid) dosing regimens would be suitable for toxicity and efficacy studies.

**Table 1:** Fundamental PK parameters of **1** after 5 mg/kg IV administration.

| **t<sub>1/2</sub>**<br>**(h)** | **C<sub>0</sub>**<br>**(ng/mL)** | **AUC<sub>last</sub>**<br>**(h\*ng/mL)** | **AUC<sub>Inf</sub>**<br>**(h\*ng/mL)** | **AUC/D**<br>**(h\*kg\*ng/**<br>**mL/mg)** | **AUC**<br>**Extr**<br>**(%)** | **MRT**<br>**(h)** | **V<sub>ss</sub>**<br>**(L/kg)** | **CL**<br>**(mL/hr/kg)** |
| --- | --- | --- | --- | --- | --- | --- | --- | --- |
| 4.35 | 33657 | 284562 | 290402 | 58080 | 2.01 | 5.83 | 0.10 | 17.4 |

Toxicity and Maximum Tolerated Dose (MTD) were assessed in neutropenic female BALB/c mice (n = 3 per dose) (**Table S3**). **1** (25, 50, and 100 mg/kg) was administered IV BID with a 12-hour interval (q12h). Animals were monitored for 5 minutes following each dose for the presence of acute toxic symptoms and autonomic effects (**Table S4**), and body weights were recorded pre-dose and 24 hours following the first treatment (**Table S5**). **1** at 100 mg/kg was fatal in the first mouse, therefore the other two mice were not administered this dose and toxicity was determined at 50 and 25 mg/kg. At 25 mg/mL, **1** did not induce mortality in any mice and no body weight drop was observed. Slight to moderate adverse effects were observed in some mice relative to the vehicle control, including increased respiration rate, tremor, and decreased spontaneous activity. No severe effects were observed relative to vehicle controls, and this dose is considered tolerated. **1** at 50 mg/mL did not induce mortality in any mice, but slight body weight losses were observed 24 hours post-treatment. Slight to moderate adverse effects including vocalization, ataxia, and hunch back were observed, indicating reduced tolerance at this dose. A full tabulation of adverse effects following each dose can be found in **Table S6** and **Table S7**. The MTD for murine efficacy models was determined to be 25 mg/kg.

*A. baumannii* ATCC 17978 was used as the infective agent in a thigh infection model to determine compound efficacy. Neutropenic BALB/c mice were inoculated intramuscularly (IM) with *A. baumannii* ATCC 17978 at 1.9 × 10^5^ CFU/mouse. At 2 h after infection, mice were treated with vehicle (10 mL/kg; IV, BID, q12h), colistin (30 mg/kg; SC, BID, q12h), or **1** (6.25, 12.5, and 25 mg/kg; IV, BID, q12h). Another infected but untreated control group was sacrificed at 2 h post infection for baseline assessment. All treatment groups were sacrificed at 26 h post infection, the thigh muscles were excised and homogenized for bacterial enumeration, and CFU/thigh was used to determine compound efficacy.

The 2 h baseline assessment group yielded bacterial counts of 5.45 log(CFU/thigh), and vehicle treated mice were at 7.80 log(CFU/thigh). Treatment with the reference agent colistin resulted in a significant decrease in bacterial load to 3.52 log(CFU/thigh). Treatment with **1** at 25 and 12.5 mg/kg yielded significant reductions in bacterial counts to 3.82 and 5.55 log(CFU/thigh), representing >99% and >98% reduction in bacterial counts relative to the vehicle control. Compared to the 2 h baseline counts, 25 mg/kg of **1** shows a significant 1.65-log_10_ reduction (p < 0.05) in bacterial counts. The 12.5 mg/kg BID dosing group shows a net bacteriostatic effect, with no increase or decrease in counts relative to the 2 h baseline. The 6.25 mg/kg dosing group did not show significant reductions in bacterial counts compared to vehicle control or 2 h baseline groups. Final results for each mouse in the efficacy study are summarized in **Table S8**.

**Figure 1:**
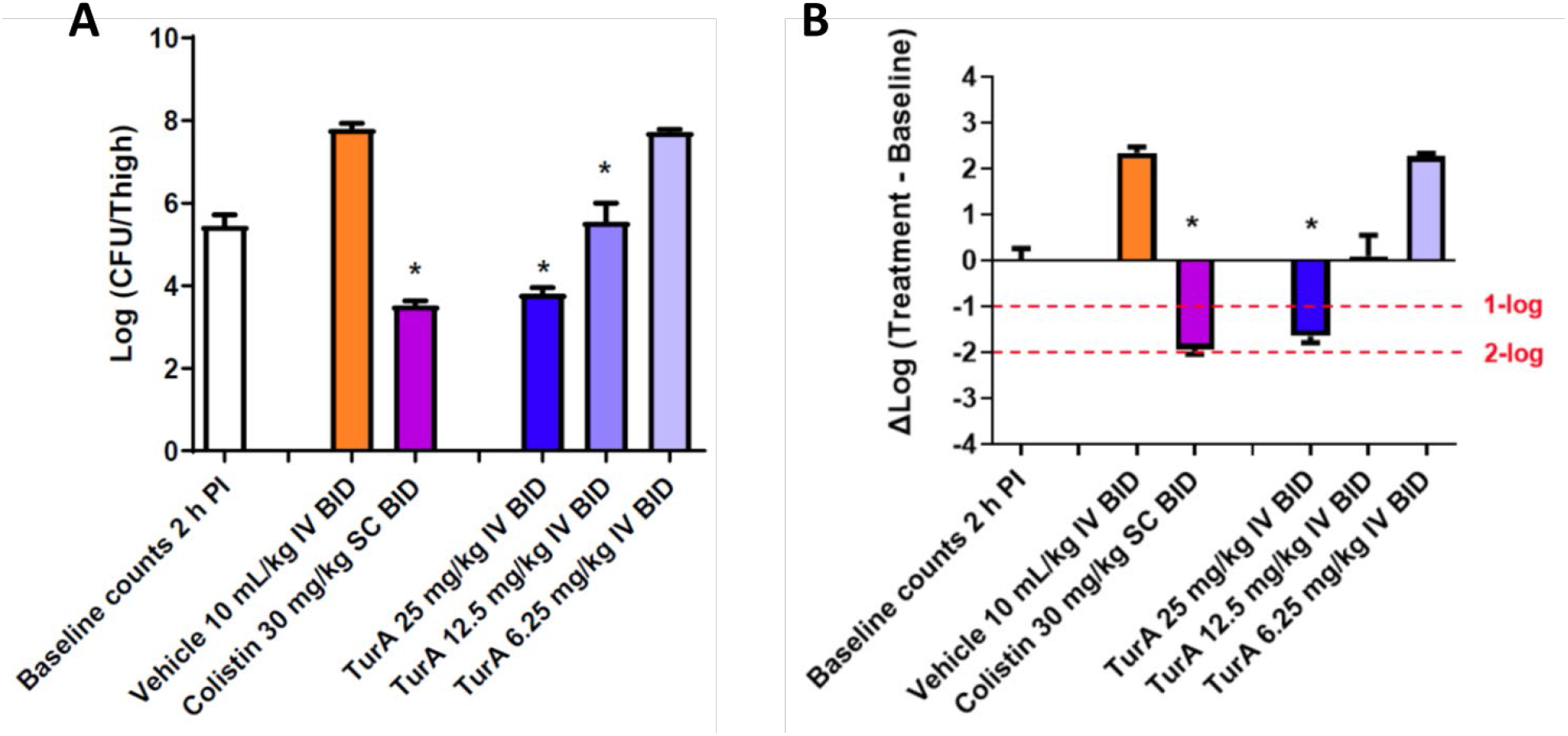
Effects of Turnercyclamycin A and colistin in the *A. baumannii* ATCC 17978 thigh infection model with neutropenic BALB/c female mice. A) Bacterial counts from excised thigh tissue for each treatment group. (*) indicates significant difference (*p* < 0.05) compared to the respective vehicle control was determined by one-way ANOVA followed by Dunnett’s test. B) Change in bacterial counts in thigh tissue at the 26 h sacrifice time point relative to the initial 2 h counts at the time of dosing. (*) indicates greater than 1-log_10_ reduction in counts relative to the baseline at the time of the first dose administration, 2 h after infection, with a significant difference (*p* < 0.05) based on one-way ANOVA followed by Dunnett’s test.

### *in vitro* resistance study

The *mcr-1* expressing plasmid was obtained from AddGene and transformed into two different strains of *Escherichia coli*: DH5α and C600. *E. coli* was used in place of Acinetobacter because both colistin and turnercyclamycins are very effective at killing E. coli, because *mcr-1* is reported to work by the same mechanism to provide *E. coli* with resistance against colistin, and because of the ease and relative safety of working with *E. coli*. The wild-type and *mcr-1* strains were treated with varying doses of colistin, turnercyclamycin A, and turnercyclamycin B. All experiments were performed in triplicate to obtain statistical significance.

Colistin fairly potently killed *E. coli*, with an MIC of 0.5 µg/mL against both wild-type *E. coli* strains, but when tested against *mcr-1* strains it was 16-fold less potent, with an MIC of 8 µg/mL. Both turnercyclamycins exhibited an MIC of 2 µg/mL against the wild-type strains, which was unchanged in 3 of 4 conditions using *mcr-1* strains. Turnercyclamycin A exhibited an MIC of 4 µg/mL against *E. coli* C600 *mcr-1*, representing a 2-fold reduction (**Table 2**). This reduction was too small to be sure of its significance. However, it paralleled a similar observation in previous work in which we used clinical isolates of colistin-resistant *Acinetobacter*. In those experiments, turnercyclamycins were usually equally effective (if not more effective) against colistin-resistant strains than wild-type strains, but in a few cases turnercyclamycin A (but not B) exhibited a 2-fold loss of potency in colistin-resistant strains. Thus, although this change is within the normal MIC error observed in experiments, it is possible that turnercyclamycin B might be more effective than turnercyclamycin A against multidrug- and colistin-resistant bacteria.

**Table 2:** Effect of *mcr-1* resistance gene on colistin and turnercyclamycin susceptibility in *E. coli* DH5α and C600

| Compound | MIC ($\mu\text{g/mL}$) | | | | | |
| --- | --- | --- | --- | --- | --- | --- |
| | DH5α | pBR322<br>in DH5α | pGDP2-MCR-1<br>in DH5α | C600 | pBR322<br>in C600 | pGDP2-MCR-1<br>in C600 |
| **colistin** | 0.5 | 0.5 | 8 | 0.5 | 1 | 8 |
| **turnercyclamycin A** | 2 | 2 | 2 | 2 | 4 | 4 |
| **turnercyclamycin B** | 2 | 2 | 4 | 4 | 4 | 4 |

## Conclusions

Here, we provide an update on the use of turnercyclamycins as antibacterial agents. Turnercyclamycin A demonstrates reasonable *in vivo* PK parameters in an IV mouse model, which are comparable to other lipopeptide antibiotics currently used in the clinic. The low volume of distribution (Vss = 0.1 L/kg) indicates that **1** does not enter cells, and thus is concentrated in the blood plasma. This is similar to the lipopeptide daptomycin, which has been shown to have high levels of protein binding in the plasma that limits its distribution. This is not an implement to efficacy, however, for bacterial infections that are extracellular, such as *Acinetorbacter*. Furthermore, the 17.4 mL/hr/kg clearance rate of the drug leads to a half-life of 4.35 h, which is within the range for a twice-daily administered antibiotic, and is an excellent starting point for optimization.

**1** was well tolerated in animals up to 25 mg/kg, with only minor and inconsistent adverse reactions observed. At this same dose, the drug was shown to be effective against *Acinetobacter baumannii* ATCC 17978 in a neutropenic mouse thigh infection model, decreasing bacterial counts from thigh tissue by 98% and lowering counts by 1.78 log10 fold compared to the baseline numbers. This demonstrates that the compound is bactericidal at these concentrations, and not simply bacteriostatic. At 2-fold lower concentrations, bacterial counts are lowered by 80%, and there is no change from the baseline counts, indicating a net bacteriostatic effect. These data are very promising, clearly demonstrating *in vivo* efficacy within a tolerated dosing regimen.

As previously reported, turnercyclamycins retain their potency against colistin-resistant clinical *Acinetobacter* complex strains. This indicates a separate molecular mechanism of action, however the exact resistance mechanisms in each clinical isolate are not fully understood. The mcr-1 resistance gene has recently threatened the use of polymyxin drugs as last line therapeutics for a number of Gram-negative infections, including *E. coli* and *Acinetobacter*. The data herein show that turnercyclamycin activity is not significantly affected by the mcr-1 plasmid, further highlighting their molecular differentiation from polymyxin-like drugs and their potential as antibiotic agents for multidrug-resistant Gram-negative infections.

## Materials

### Animals

#### PK and MTD studies

BALB/c female mice, weighing 18 ± 2 g, were provided by BioLasco Taiwan (under Charles River Laboratories Licensee). All animals were maintained in a well-controlled temperature (20 - 24°C) and humidity (30% - 70%) environment with 12 hours light/dark cycles. Free access to standard lab diet [MFG (Oriental Yeast Co., Ltd., Japan)] and autoclaved tap water were granted. All aspects of this work including housing, experimentation, and animal disposal were performed in general accordance with the “Guide for the Care and Use of Laboratory Animals: Eighth Edition*”* (National Academies Press, Washington, D.C., 2011) in our AAALAC-accredited laboratory animal facility. In addition, the animal care and use protocol was reviewed and approved by the IACUC at Pharmacology Discovery Services Taiwan, Ltd).

#### Efficacy study

BALB/c female mice, 7 weeks of age, were provided by BioLasco Taiwan (under Charles River Laboratories Licensee). Animals were acclimated for 3 days prior to use and were confirmed to be in good health. The space allocation for animals was 39 × 20 × 16 cm. All animals were maintained in a hygienic environment with controlled temperature (20 - 24°C), humidity (30% - 70%) and 12 hours light/dark cycles. Free access to sterilized standard lab diet [MFG (Oriental Yeast Co., Ltd., Japan)] and autoclaved tap water were granted for study period. All aspects of this work including housing, experimentation, and animal disposal were performed in general accordance with the “Guide for the Care and Use of Laboratory Animals: Eighth Edition” (National Academies Press, Washington, D.C., 2011). The study was performed in our AAALAC-accredited ABSL-2 laboratory animal facility under the supervision of veterinarians. In addition, the animal care and use protocol was reviewed and approved by the IACUC at Pharmacology Discovery Services Taiwan, Ltd.

### Organisms

*Acinetobacter baumannii* (ATCC 17978) was acquired from the American Type Culture Collection (Rockville, MD, USA). DH5α competent cells (ThermoFisher 18265017) was acquired from ThermoFisher Scientific. *E. coli* C600 strains were a kind gift from Dr. Matthew Mulvey of the Department of Pathology, University of Utah, USA. All bacterial strains are cryopreserved as single-use frozen working stock cultures stored at −80 °C.

### Equipment

0-1000 g Electronic scale (Tanita Corporation, Japan), Animal cage (Allentown, USA), Centrifuge 5810R (Eppendrof, Germany), Disposal syringe (1 mL, Terumo Corporation, Japan), Gilson Pipettes (# P200 Neo-P10N Micro Pipette; Gilson, France), ImprominiTM EDTA-K2 tube (Improve Medical, China), LC-MS/MS Triple Quad™ 5500+ (SCIEX, USA), Microcentrifuge tubes 1.5 mL click-cap (Treff AG, Switzerland), Phenomenex SynergiTM 4μm, Polar-RP 80A LC Column 50 × 2mm(Phenomenex, USA), Polypropylene 96-well round U-bottom deepwell plates and silicone microplate lids (StorPlate-96 U and StorMat-96; PerkinElmer Inc., USA), RAININ Pipettes (E4 Multi E12-50XLS+, E12-300XLS, Refurbished Rainin E4™ XLS™ electronic 12 channel pipette, 30-300 uL, E4 Pipette Multi E12-1200XLS+, and EA6-300XLS; RAININ, USA), Pipette Tips (Costar, USA), and Stop watch (Casio, China).

Absorbance microplate readers (Tecan, Infinite F50, USA), Biological safety cabinet (NuAire, USA), Centrifuge (Model 5922, Kubota, Japan), Individually ventilated cages (GM500 IVC seal safe plus cage system) (Tecniplast, Italy), Orbital shaking incubator (Firstek Scientific, Taiwan), Pipetman (Rainin, USA), Polytron homogenizer (Kinematica, Switzerland), Portable weighing scale (JKH-1000, Jadever, Taiwan), Refrigerated incubator (Hotpack, USA), and Ultra-low temperature freezer (Panasonic, Japan).

## Methods

### Plasma Sample Collection from Mice (Parallel sampling)

Blood aliquots were collected via cardiac puncture (0.3 mL). Blood aliquots were collected in tubes coated with EDTA-K2, mixed gently, then kept on ice and centrifuged at 2,500 ×g for 15 minutes at 4°C, within 1 hour of collection. The plasma (supernatant) was then harvested and kept frozen at −70°C until further processing.

### Quantitative Bioanalysis (Plasma Samples)

The plasma samples were processed using acetonitrile precipitation and analyzed by LC-MS/MS. The detailed chromatographic conditions, bioanalytical methods, and acceptance criteria are summarized as followings:

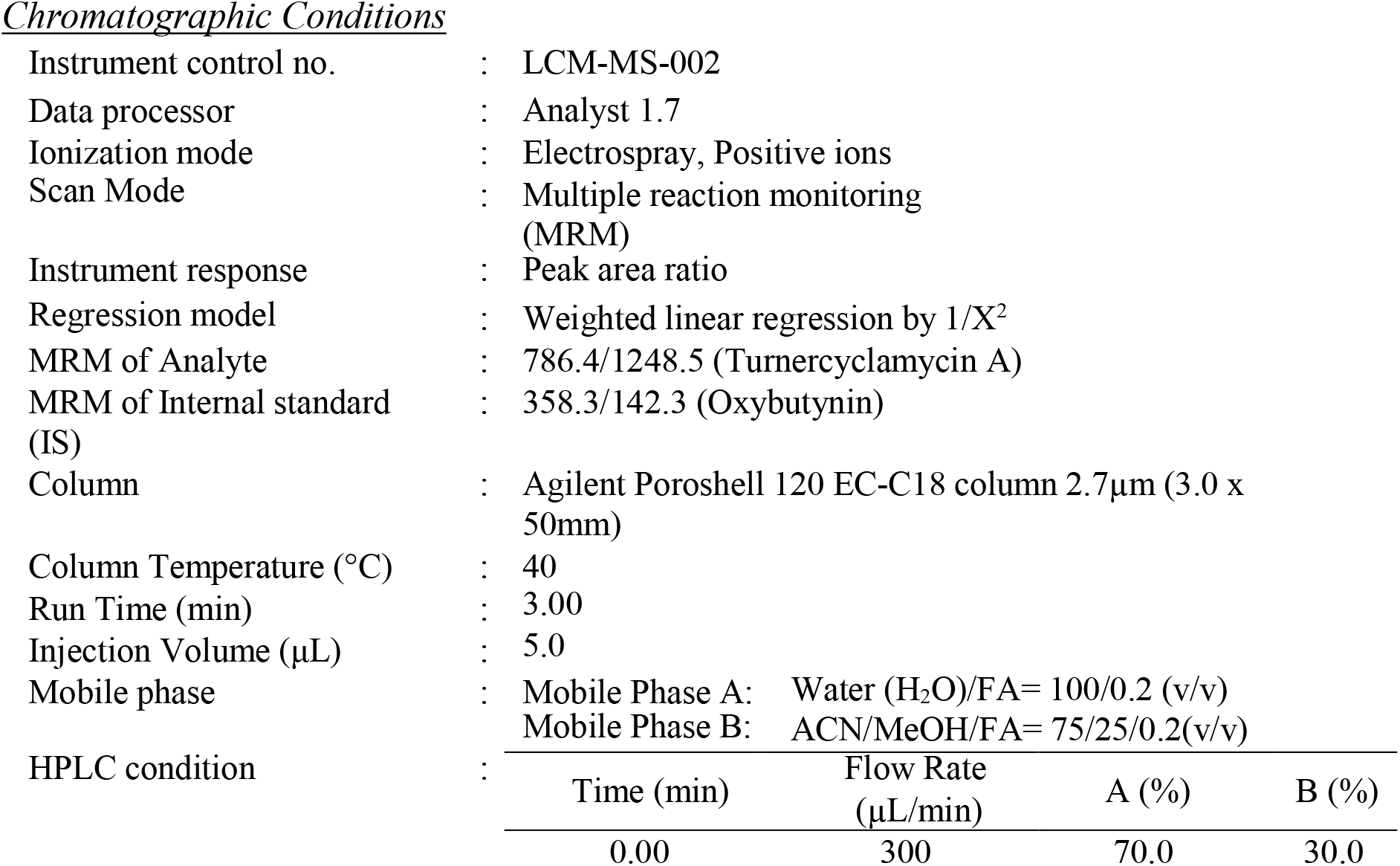
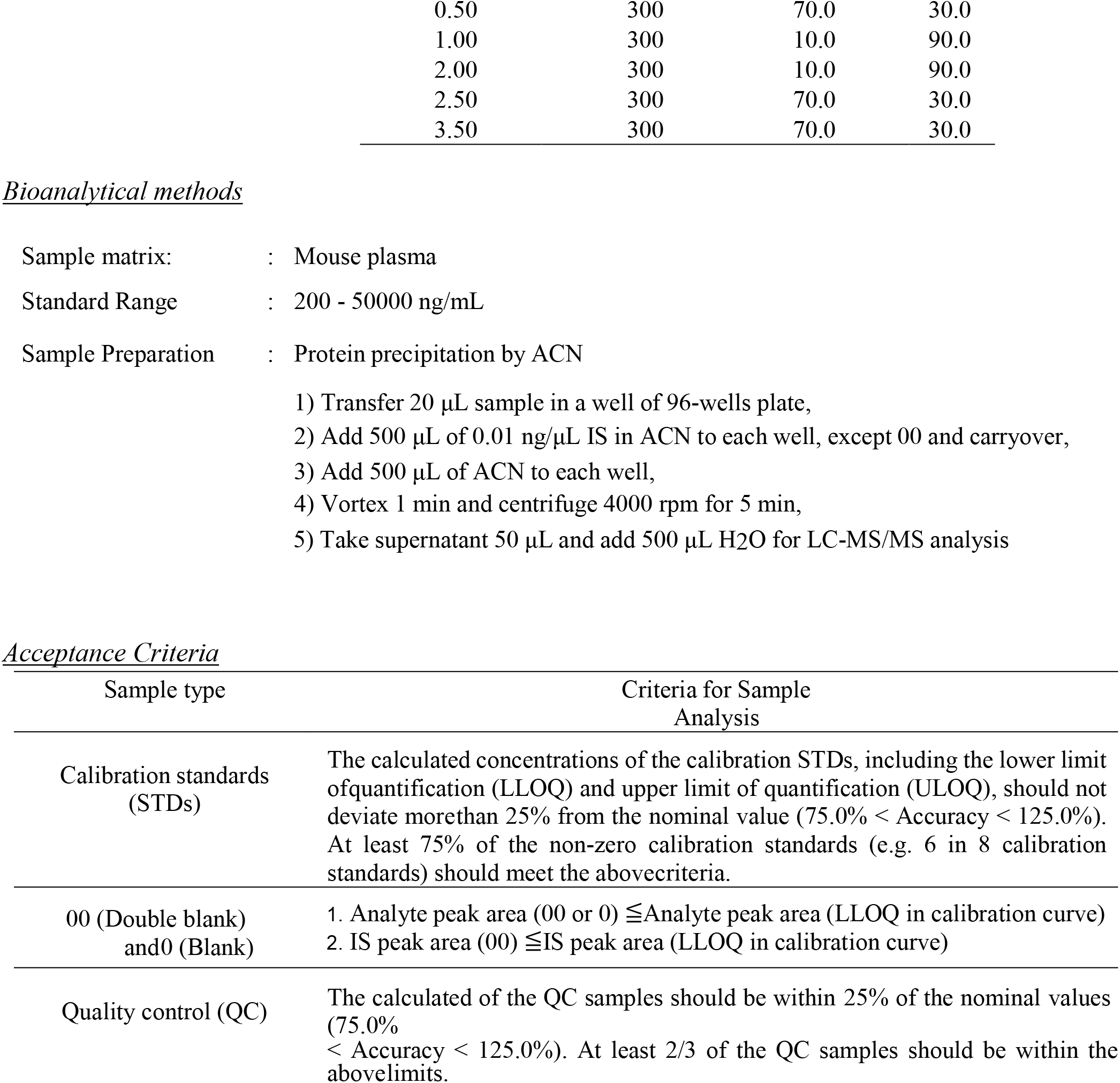

### Pharmacokinetics

The exposure levels (ng/mL) of Turnercyclamycin A in plasma samples were determined by LC-MS/MS. Plots of plasma concentrations (mean ± SD) vs. time for Turnercyclamycin A were constructed. The PK parameters of the test compound after IV (t1/2, C0, AUClast, AUCInf, AUCExtr, MRT, Vss, and CL) were obtained from the non-compartmental analysis (NCA) of the plasma data using WinNonlin.

### Antimicrobial microdilution assay

Glycerol stocks of untransformed *E. coli* DH5α and *E. coli* C600 were streaked on Luria-Bertani (LBA) without selective antibiotics. *E. coli* DH5α and *E. coli* C600 transformed with empty vector pBR322 (NEB N3033S) and pGPDP2 MCR-1 (Addgene 118404) were streaked on (LBA) with selective antibiotics, and all plates were incubated overnight at 37°C. Single colonies from each plate where then transferred to LB broth and incubated for 6-8h at 30°C, 150rpm. Turbidity was adjusted to match 0.5 McFarland standard (1×10^8^ cells/mL), then diluted 200-fold to serve as the inoculum for the assay. 200 µL of the test organism was added to each well of a 96-well plate. Compounds were prepared in neat, sterile DMSO and serially diluted using a two-fold dilution scheme, then added to each well resulting in eight dilutions for each compound starting at 32µg/mL for turnercyclamycins and 8µg/mL for colistin. The plate was then incubated for 18-20h at 30°C, 150rpm. After incubation, 10µL of 5mg/mL MTT was added to the wells and incubated for 2h. A_570_ was the measured using Biotek-Synergy 2 Microplate Reader (Biotek) and percent inhibition computed using absorbance values measured.

## References

(1) Centers for Disease Control and Prevention (U.S.). Antibiotic Resistance Threats in the United States, 2019; Centers for Disease Control and Prevention (U.S.), 2019. https://doi.org/10.15620/cdc:82532.

(2) Peleg, A. Y.; Seifert, H.; Paterson, D. L. Acinetobacter Baumannii: Emergence of a Successful Pathogen. Clinical Microbiology Reviews 2008, 21 (3), 538–582. https://doi.org/10.1128/CMR.00058-07.

(3) Falagas, M. E.; Kasiakou, S. K. Colistin: The Revival of Polymyxins for the Management of Multidrug-Resistant Gram-Negative Bacterial Infections. Clin Infect Dis 2005, 40 (9), 1333–1341. https://doi.org/10.1086/429323.

(4) Li, J.; Nation, R. L.; Turnidge, J. D.; Milne, R. W.; Coulthard, K.; Rayner, C. R.; Paterson, D. L. Colistin: The Re-Emerging Antibiotic for Multidrug-Resistant Gram-Negative Bacterial Infections. Lancet Infect Dis 2006, 6 (9), 589–601. https://doi.org/10.1016/S1473-3099(06)70580-1.

(5) Rodriguez, C. H.; Hernan, R. C.; Bombicino, K.; Karina, B.; Granados, G.; Gabriela, G.; Nastro, M.; Marcela, N.; Vay, C.; Carlos, V.; Famiglietti, A.; Angela, F. Selection of Colistin-Resistant Acinetobacter Baumannii Isolates in Postneurosurgical Meningitis in an Intensive Care Unit with High Presence of Heteroresistance to Colistin. Diagn Microbiol Infect Dis 2009, 65 (2), 188–191. https://doi.org/10.1016/j.diagmicrobio.2009.05.019.

(6) Miller, B. W.; Lim, A. L.; Lin, Z.; Bailey, J.; Aoyagi, K. L.; Fisher, M. A.; Barrows, L. R.; Manoil, C.; Schmidt, E. W.; Haygood, M. G. Shipworm Symbiosis Ecology-Guided Discovery of an Antibiotic That Kills Colistin-Resistant Acinetobacter. Cell Chemical Biology 2021. https://doi.org/10.1016/j.chembiol.2021.05.003.

