## Supplemental Figures and Tables for "Initial efficacy determination and resistance profile of anti-*Acinetobacter* antibiotics, turnercyclamycins"

#### Table of Contents

### Figures

**Figure S1**: Plasma concentration of turnercyclamycin A following 5 mg/kg injection for PK study

**Figure S2**: Representative MIC data for colistin, turnercyclamycin A, and turnercyclamycin B in untransformed, empty pBR322 transformed, and pGDP2-MCR-1 transformed *E. coli* DH5α.

**Figure S3**: Representative MIC data for colistin, turnercyclamycin A, and turnercyclamycin B in untransformed, empty pBR322 transformed, and pGDP2-MCR-1 transformed *E. coli* C600

### Tables

**Table S1**: Experimental design for PK study

**Table S2**: Exposure levels (ng/mL) of turnercyclamycin A in plasma samples in PK study

**Table S3**: Experimental design for toxicity/MTD study

**Table S4**: Clinical symptoms for observation during toxicity/MTD study

**Table S5**: Body weight of mice in toxicity/MTD study

**Table S6**: Adverse effects within 5 minutes of 1st dose in toxicity/MTD study

**Table S7**: Adverse effects within 5 minutes of 2nd dose in toxicity/MTD study

**Table S8:** Final results per mouse of thigh infection efficacy study

### Figures

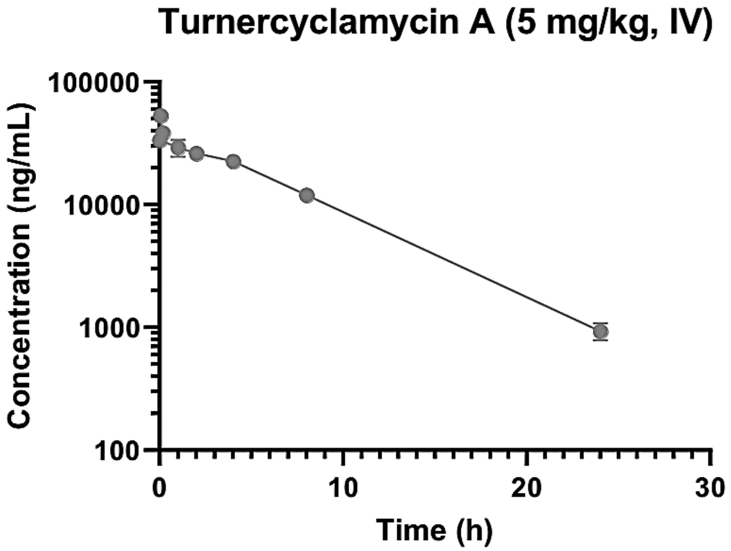


**Figure S1**: Plasma concentration of turnercyclamycin A following 5 mg/kg injection for PK study

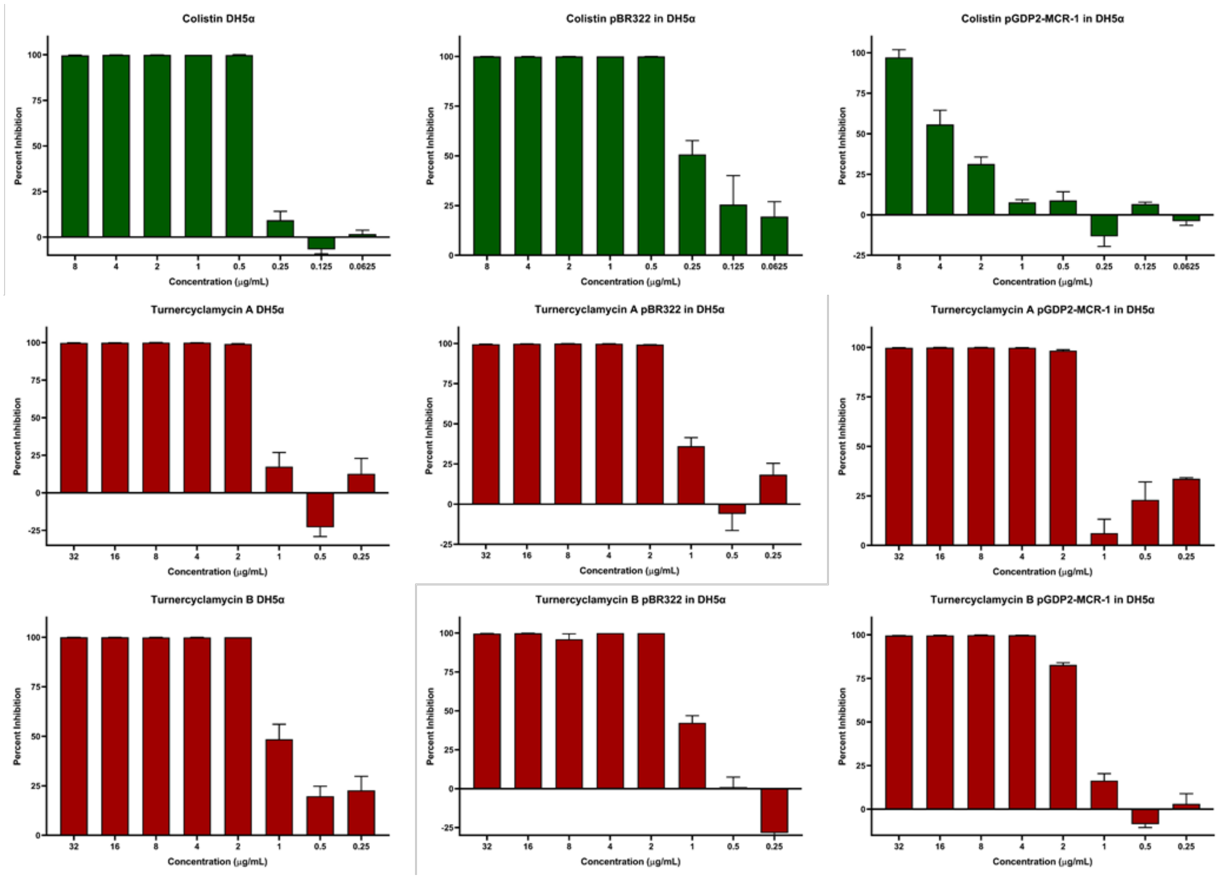


**Figure S2**: Representative MIC data for colistin, turnercyclamycin A, and turnercyclamycin B in untransformed, empty pBR322 transformed, and pGDP2-MCR-1 transformed *E. coli* DH5α.

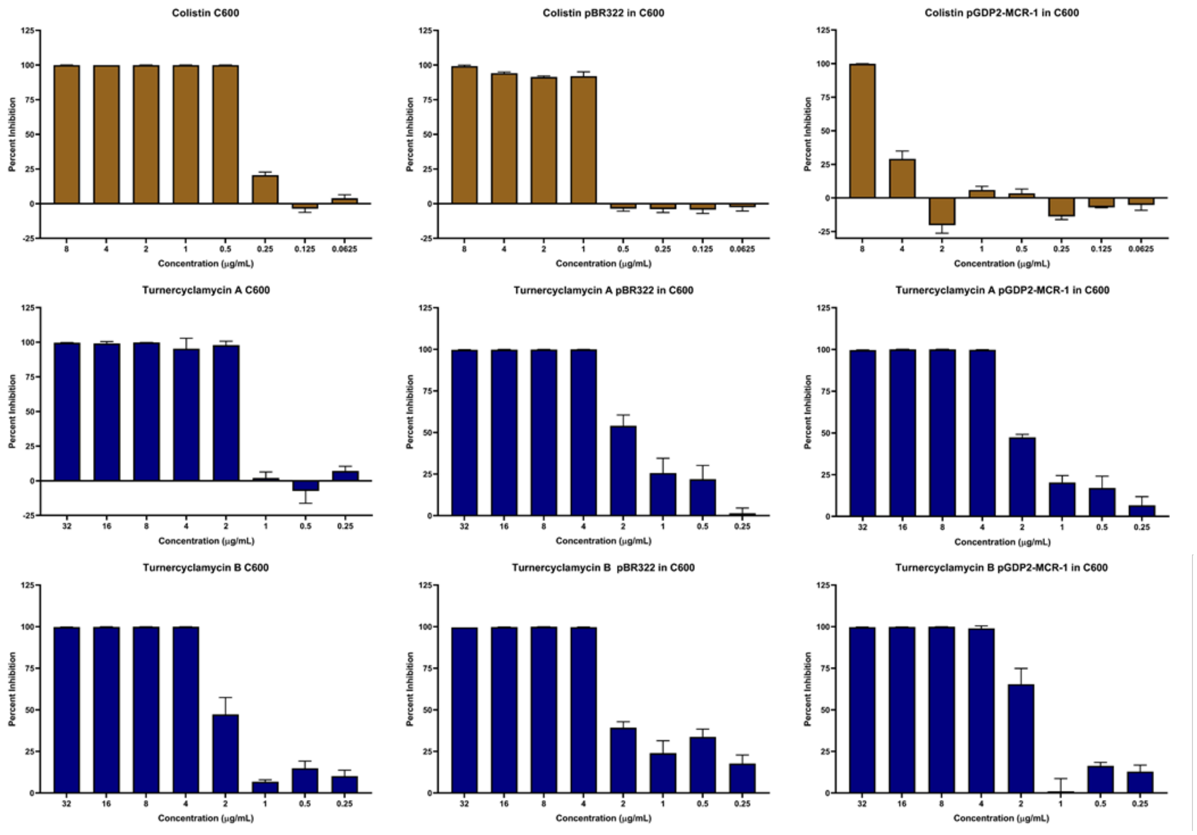


**Figure S3**: Representative MIC data for colistin, turnercyclamycin A, and turnercyclamycin B in untransformed, empty pBR322 transformed, and pGDP2-MCR-1 transformed *E. coli* C600.

### Tables

**Table S1**: Experimental design for PK study

| **Animal Use** | |
|---|---|
| Animal Species/Strain | Mouse/ Female BALB/c |
| Body Weight | 18 ± 2 g |
| Fasting Regimen | None |
| Pre-dose Observations | Body weight (gm) |
| **Test Article Dosing and PK Sample Collection** | |
| Test Article ID | Turnercyclamycin A |
| Route(s) of Administration | IV |
| Dose Levels (mg/kg) | 5 |
| Dose Volume (mL/kg) | 5 |
| Proposed Formulation/Vehicle | 3% ethanol/ PBS |
| Number of Animals per Dosing Group | 24 |
| Total Number of Animals | 24 |
| Total Number of Samples | 24 plasma |
| PK Sample (Plasma) Collection Time | 0.05, 0.167, 0.5, 1, 2, 4, 8, and 24 hour(s) |
| Target Blood Sample Volume (mL) | 0.3 mL via cardiac puncture |
| Preferred Anticoagulation | EDTA-$K_2$ |
| Preferred Sample Storage | $\leq$-70 °C |

**Table S2**: Exposure levels (ng/mL) of turnercyclamycin A in plasma samples in PK study

| Time (h) | Sample concentration (ng/mL) | | | Mean | Standard Deviation |
|---|---|---|---|---|---|
| 0.05 | 52235 | 51887 | 55877 | 53333 | 2210 |
| 0.167 | 36107 | 38906 | 40924 | 38646 | 2419 |
| 0 | 34373 | 31954 | 34645 | 33657 | 1481 |
| 1 | 24635 | 33834 | 29337 | 29269 | 4600 |
| 2 | 26373 | 25446 | 26782 | 26200 | 685 |
| 4 | 22923 | 22209 | 22495 | 22542 | 359 |
| 8 | 11661 | 13130 | 11123 | 11971 | 1039 |
| 24 | 974 | 764 | 1054 | 931 | 150 |

**Table S3**: Experimental design for toxicity/MTD study

| Group | Test Substance | Dose Schedule | Dose Route | Conc. mg/mL | Dosage | | | Mouse BALB/c (female)[d] |
|---|---|---|---|---|---|---|---|---|
| | | | | | mL/kg | mg/kg/dose | mg/kg/24h | |
| 1 | Vehicle[a] | bid (q12h)[b] | IV[c] | NA | 10 | NA | NA | 3 |
| 2 | Turnercyclamycin A (C. factor: 1.010) | bid (q12h)[b] | IV[c] | 10 | 10 | 100 | 200 | 3 |
| 3 | Turnercyclamycin A (C. factor: 1.010) | bid (q12h)[b] | IV[c] | 5 | 10 | 50 | 100 | 3 |
| 4 | Turnercyclamycin A (C. factor: 1.010) | bid (q12h)[b] | IV[c] | 2.5 | 10 | 25 | 50 | 3 |

[a] Vehicle: 3% ethanol/ PBS
[b] administered twice per day (BID) at 12 hour intervals (q12h) for 24 h
[c] IV bolus in slow injection at least for 10 secounds
[d] Neutropenic BALB/c mice were used for this study

**Table S4**: Clinical symptoms for observation during toxicity/MTD study

| Body Weight (B.W.) (g) | Decrease in Touch Response | Decrease in Spontaneous Activity | Low Limb Post | Body Temperature |
|---|---|---|---|---|
| Irritability | Increase in Exploration | Straub Tail | Skin Color | Piloerection |
| Hyperactivity | Decrease in Exploration | Reactivity | Respiration | Increase in Palpebral Size |
| Increase in Startle Response | Pinna | Righting | Salivation (Fluid and Viscosity) | Decrease in Palpebral Size |
| Increase Touch Response | Placing | Ataxia | Lacrimation | Death |
| Decrease Startle Response | Tremor | Convulsion (Chronic / Tonic) | Diarrhea | |

**Table S5**: Body weight of mice in toxicity/MTD study

| Compound | Route | Dose (mg/kg) | No. | Gender | Body Weight ( g ) | | | | |
|---|---|---|---|---|---|---|---|---|---|
| | | | | | Day -4 | Day -1 | Day 0 (first) | Day 0 (second) | Day 1 |
| Vehicle (3% ethanol/ PBS) | IV | 10 mL/kg bid (q12h) x1 | 1 | Female | 16 | 16 | 16 | 15 | 16 |
| | | | 2 | | 16 | 17 | 17 | 16 | 16 |
| | | | 3 | | 16 | 16 | 17 | 16 | 16 |
| PT# 1250365 (UUM-1) Turnercyclamycin A | IV | 100 [a] bid (q12h) x1 | 1 | Female | 17 | 16 | 15 | NA | |
| | | | 2 | | 17 | 16 | 16 | | |
| | | | 3 | | 17 | 17 | 16 | | |
| | IV | 50 bid (q12h) x1 | 4 | Female | 17 | 16 | 16 | 15 | 15 |
| | | | 5 | | 17 | 16 | 15 | 15 | 14 |
| | | | 6 | | 16 | 17 | 16 | 16 | 16 |
| | IV | 25 bid (q12h) x1 | 7 | Female | 18 | 17 | 17 | 16 | 17 |
| | | | 8 | | 17 | 17 | 17 | 16 | 17 |
| | | | 9 | | 17 | 17 | 17 | 17 | 17 |

**Table S6**: Adverse effects within 5 minutes of 1st dose in toxicity/MTD study

| Treatment | Vehicle (3% ethanol/ PBS) | | | PT# 1250365 (UUM-1) Turnercyclamycin A | | | | | | | | |
|---|---|---|---|---|---|---|---|---|---|---|---|---|
| Route | IV, bid (q12h) x 1 | | | | | | | | | | | |
| Dosage | 10 mL/kg | | | 100 mg/kg a | | | 50 mg/kg | | | 25 mg/kg | | |
| Observation time | 5 minutes after the 1st dose | | | | | | | | | | | |
| Gender | Female | | | | | | | | | | | |
| BEHAVIORAL | No. 1 | No. 2 | No. 3 | No. 1 | No. 2 | No. 3 | No. 1 | No. 2 | No. 3 | No. 1 | No. 2 | No. 3 |
| B.W. (g) | - | - | - | | | | - | - | - | - | - | - |
| Irritability | - | - | - | | | | - | - | - | - | - | - |
| Hyperactivity | - | - | - | | | | - | - | - | - | - | - |
| Inc. Startle | - | - | - | | | | - | - | - | - | - | - |
| Inc. Touch | - | - | - | | | | - | - | - | - | - | - |
| Dec. Startle Response | - | - | - | | | | - | - | - | - | - | - |
| Dec. Touch Response | - | - | - | | | | - | - | - | - | - | - |
| Inc. Exploration | - | - | - | | | | - | - | - | - | - | - |
| Dec. Exploration | - | - | - | | | | - | - | - | - | - | - |
| Pinna | - | - | - | | | | - | - | - | - | - | - |
| Placing | - | - | - | | | | - | - | - | - | - | - |
| NEUROLOGIC | | | | | | | | | | | | |
| Tremor | - | - | - | | | | - | - | ± | - | ± | - |
| Dec. Spont. Activity | - | - | - | | | | ± | - | - | - | - | - |
| Straub Tail | - | - | - | | | | - | - | - | - | - | - |
| Reactivity | - | - | - | | | | ± | - | ± | - | - | - |
| Righting | - | - | - | | | | - | - | - | - | - | - |
| Ataxia | - | - | - | | | | - | - | - | - | - | - |
| Convulsion C.T.C-T | - | - | - | | | | - | - | - | - | - | - |
| Low Limb Post | + | ± | - | | | | ± | - | ± | - | ± | + |
| Abdominal Tone | + | + | + | | | | + | + | + | + | + | ± |
| Limb Tone | + | + | + | | | | + | + | + | + | + | + |
| Grip Strength | - | - | - | | | | - | - | - | - | - | - |
| AUTONOMIC | | | | | | | | | | | | |
| Skin Color | - | - | - | | | | - | - | - | - | - | - |
| Respiration | - | - | - | | | | Fast ± | Fast ± | - | - | - | - |
| Salivation F.V. | - | - | - | | | | - | - | - | - | - | - |
| Lacrimation | - | - | - | | | | - | - | - | - | - | - |
| Diarrhea | - | - | - | | | | - | - | - | - | - | - |
| Body Temperature | - | - | - | | | | - | - | - | - | - | - |
| Piloerection | - | ± | ± | | | | ± | ± | ± | ± | ± | ± |
| Inc. Palpebral Size | - | - | - | | | | - | - | - | - | - | - |
| Dec. Palpebral Size | - | - | - | | | | - | - | - | - | - | - |
| Others | - | - | - | | | | - | - | - | - | - | - |
| Death | - | - | - | + | | | - | - | - | - | - | - |

**Table S7**: Adverse effects within 5 minutes of 2nd dose in toxicity/MTD study

| Treatment | Vehicle (3% ethanol/ PBS) | | | PT# 1250365 (UUM-1) Turnercyclamycin A | | | | | | | | |
|---|---|---|---|---|---|---|---|---|---|---|---|---|
| Route | IV, bid (q12h) x 1 | | | | | | | | | | | |
| Dosage | 10 mL/kg | | | 100 mg/kg a | | | 50 mg/kg | | | 25 mg/kg | | |
| Observation time | 5 minutes after the 2nd dose | | | | | | | | | | | |
| Gender | Female | | | | | | | | | | | |
| BEHAVIORAL | No. 1 | No. 2 | No. 3 | No. 1 | No. 2 | No. 3 | No. 1 | No. 2 | No. 3 | No. 1 | No. 2 | No. 3 |
| B.W. (g) | - | - | - | | | | - | - | - | - | - | - |
| Irritability | - | - | - | | | | Voc | Voc | - | Voc | - | - |
| Hyperactivity | - | - | - | | | | - | - | - | - | - | - |
| Inc. Startle | - | - | - | | | | - | - | - | - | - | - |
| Inc. Touch | - | - | - | | | | ± | - | - | - | - | - |
| Dec. Startle Response | - | - | - | | | | - | - | - | - | - | - |
| Dec. Touch Response | - | - | - | | | | - | - | - | - | - | - |
| Inc. Exploration | - | - | - | | | | - | - | - | - | - | - |
| Dec. Exploration | - | - | - | | | | - | - | - | - | - | - |
| Pinna | - | - | - | | | | - | - | - | - | - | - |
| Placing | - | - | - | | | | - | - | - | - | - | - |
| NEUROLOGIC | | | | | | | | | | | | |
| Tremor | - | - | - | | | | - | - | - | - | - | - |
| Dec. Spont. Activity | - | - | - | | | | ± | - | ± | - | - | ± |
| Straub Tail | - | - | - | | | | - | - | - | - | - | - |
| Reactivity | - | - | - | | | | - | - | ± | - | - | - |
| Righting | - | - | - | | | | - | - | - | - | - | - |
| Ataxia | - | - | - | | | | ± | - | ± | - | - | - |
| Convulsion C.T.C-T | - | - | - | | | | - | - | - | - | - | - |
| Low Limb Post | + | - | - | | | | + | + | + | + | ± | ± |
| Abdominal Tone | + | + | + | | | | + | + | + | + | + | + |
| Limb Tone | + | + | + | | | | + | + | + | + | + | + |
| Grip Strength | - | - | - | | | | - | - | - | - | - | - |
| AUTONOMIC | | | | | | | | | | | | |
| Skin Color | - | - | - | | | | - | - | - | - | - | - |
| Respiration | - | Fast ± | Fast ± | | | | Fast ± | Fast ± | Fast ± | - | Fast ± | Fast ± |
| Salivation F.V. | - | - | - | | | | - | - | - | - | - | - |
| Lacrimation | - | - | - | | | | - | - | - | - | - | - |
| Diarrhea | - | - | - | | | | - | - | - | - | - | - |
| Body Temperature | - | - | - | | | | - | - | - | - | - | - |
| Piloerection | - | ± | ± | | | | ± | ± | ± | ± | ± | ± |
| Inc. Palpebral Size | - | - | - | | | | - | - | - | - | - | - |
| Dec. Palpebral Size | - | - | - | | | | - | - | - | - | - | - |
| Others | - | - | - | | | | H.B. | H.B. | H.B. | - | - | - |
| Death | - | - | - | | | | - | - | - | - | - | - |

**Table S8:** Final results per mouse of thigh infection efficacy study

| Group | Treatment | Dose Route / Schedule | Animal No. | Thigh Weight (g) | CFU/thigh Time post inoculation 2, 26 h | Decrease (%) | Log (CFU/thigh) Time post inoculation 2, 26 h | Δ |
|---|---|---|---|---|---|---|---|---|
| 1 | Baseline, 2 h post infection | N/A | 1 | 0.741 | $4.20 \times 10^5$ | | 5.62 | |
| | | | 2 | 0.683 | $6.99 \times 10^5$ | | 5.84 | |
| | | | 3 | 0.766 | $4.14 \times 10^5$ | | 5.62 | |
| | | | 4 | 0.798 | $5.91 \times 10^5$ | | 5.77 | |
| | | | 5 | 0.748 | $2.64 \times 10^4$ | | 4.42 | |
| | | | Mean | 0.747 | $4.30 \times 10^5$ | -- | 5.45 | -- |
| | | | SEM | 0.019 | $1.14 \times 10^5$ | | 0.26 | |
| 2 | Vehicle (3% ethanol/ PBS) | 10 mL/kg IV, BID, q12h | 1 | 0.725 | $1.21 \times 10^8$ | | 8.08 | |
| | | | 2 | 0.738 | $2.96 \times 10^7$ | | 7.47 | |
| | | | 3 | 0.826 | $4.10 \times 10^7$ | | 7.61 | |
| | | | 4 | 0.813 | $1.30 \times 10^8$ | | 8.11 | |
| | | | 5 | 0.751 | $5.40 \times 10^7$ | | 7.73 | |
| | | | Mean | 0.771 | $7.51 \times 10^7$ | -- | 7.80 | 2.35 |
| | | | SEM | 0.020 | $2.10 \times 10^7$ | | 0.13 | |
| 3 | Colistin | 30 mg/kg SC, BID, q12h | 1 | 0.718 | $1.77 \times 10^3$ | | 3.25 | |
| | | | 2 | 0.652 | $5.73 \times 10^3$ | | 3.76 | |
| | | | 3 | 0.648 | $5.64 \times 10^3$ | | 3.75 | |
| | | | 4 | 0.644 | $1.98 \times 10^3$ | | 3.30 | |
| | | | 5 | 0.739 | $3.57 \times 10^3$ | | 3.55 | |
| | | | Mean | 0.680 | $3.74 \times 10^3$ | 100 | 3.52* | -2.73[#] |
| | | | SEM | 0.020 | $8.54 \times 10^2$ | | 0.11 | |
| 4 | PT# 1251595 (UUM-2) (Turnercyclamycin A) | 25 mg/kg IV, BID, q12h | 1 | 0.644 | $5.91 \times 10^3$ | | 3.77 | |
| | | | 2 | 0.691 | $7.68 \times 10^3$ | | 3.89 | |
| | | | 3 | 0.602 | $8.76 \times 10^3$ | | 3.94 | |
| | | | 4 | 0.713 | $1.38 \times 10^4$ | | 4.14 | |
| | | | 5 | 0.709 | $1.89 \times 10^3$ | | 3.28 | |
| | | | Mean | 0.672 | $7.61 \times 10^3$ | 100 | 3.80* | -1.65[#] |
| | | | SEM | 0.021 | $1.94 \times 10^3$ | | 0.14 | |
| 5 | PT# 1251595 (UUM-2) (Turnercyclamycin A) | 12.5 mg/kg IV, BID, q12h | 1 | 0.783 | $5.67 \times 10^5$ | | 5.75 | |
| | | | 2 | 0.803 | $7.02 \times 10^6$ | | 6.85 | |
| | | | 3 | 0.771 | $7.89 \times 10^5$ | | 5.90 | |
| | | | 4 | 0.766 | $1.32 \times 10^5$ | | 5.12 | |
| | | | 5 | 0.752 | $1.35 \times 10^4$ | | 4.13 | |

| Group | Treatment | Dose Route / Schedule | Animal No. | Thigh Weight (g) | CFU/thigh Time post inoculation 2, 26 h | Decrease (%) | Log (CFU/thigh) Time post inoculation 2, 26 h | Δ |
|---|---|---|---|---|---|---|---|---|
| | | | Mean | 0.775 | $1.70 \times 10^6$ | 98 | 5.55* | 0.10 |
| | | | SEM | 0.009 | $1.34 \times 10^6$ | | 0.45 | |
| 6 | PT# 1251595 (UUM-2) (Turnercyclamycin A) | 6.25 mg/kg IV, BID, q12h | 1 | 0.782 | $4.59 \times 10^7$ | | 7.66 | |
| | | | 2 | 0.821 | $3.63 \times 10^7$ | | 7.56 | |
| | | | 3 | 0.788 | $4.62 \times 10^7$ | | 7.66 | |
| | | | 4 | 0.753 | $6.93 \times 10^7$ | | 7.84 | |
| | | | 5 | 0.000 | $7.71 \times 10^7$ | | 7.89 | |
| | | | Mean | 0.629 | $5.50 \times 10^7$ | 27 | 7.72 | 2.27 |
| | | | SEM | 0.158 | $7.76 \times 10^6$ | | 0.06 | |
